# Freezing and water availability structure the evolutionary diversity of trees across the Americas

**DOI:** 10.1101/728717

**Authors:** R. A. Segovia, R. T. Pennington, T. R. Baker, F. Coelho de Souza, D. M. Neves, C. C. Davis, J. J. Armesto, A. T. Olivera-Filho, K. G. Dexter

## Abstract

The historical course of evolutionary diversification shapes the current distribution of biodiversity, but the main forces constraining diversification are unclear. We unveil the evolutionary structure of tree species diversity across the Americas to assess whether an inability to move (dispersal limitation) or to evolve (niche conservatism) is the predominant constraint in plant diversification and biogeography. We find a fundamental divide in tree lineage composition between tropical and extratropical environments, defined by the absence versus presence of freezing temperatures. Within the Neotropics, we uncover a further evolutionary split between moist and dry forests. Our results demonstrate that American tree lineages, though broadly distributed geographically, tend to retain their ancestral environmental relationships and that phylogenetic niche conservatism is the primary force structuring the distribution of tree biodiversity.

## Main text

A central challenge in biogeography and macroevolution is to understand the primary forces that drove the diversification of life. Was diversification confined within continents, and characterized by adaptation of lineages to different major environments (i.e., biome switching), or did lineages tend to disperse across great distances, but retain their ancestral environmental niche (i.e., phylogenetic niche conservatism)? Classically, the attempts to define biogeographic regions based on shared plant and animal distributions lend support to the first hypothesis, that large-scale patterns may be explained by regionally confined evolutionary diversification, rather than long-distance dispersal (*1–3*). However, recent studies of the distribution of plant lineages at global scales have documented high levels of inter-continental dispersal (e.g., *4–8*), and revealed that lineages tend to retain their ancestral biomes when dispersing (*9, 10*). These latter findings suggest that environmental associations of lineages may be the primary force organizing the course of diversification, but we lack studies comparing the degree of evolutionary similarity between species assemblages at broad scales to adequately resolve this debate.

With high mountain chains running north to south across latitudes and a mosaic of contrasting environments, the Americas represent a natural laboratory to investigate the evolutionary forces behind the modern distribution of biodiversity. Here, we examine the phylogenetic composition of angiosperm tree assemblages across the Americas as a means to determine whether dispersal limitation or phylogenetic niche conservatism had a greater impact on the present-day evolutionary structure of biodiversity. If lineages tend to retain their environmental niche as they diversify across space, we would expect major evolutionary groups to be restricted to specific environmental regimes. This leads to the prediction that lineage composition of assemblages from extratropical regions in both hemispheres should be more similar to each other than to assemblages occurring in intervening tropical regions. In addition, we would predict that assemblages from dry tropical environments should show greater similarity in tree lineage composition to each other than to assemblages from moist environments with which they may be spatially contiguous (*11*). Alternatively, if diversification is spatially restricted and biome switching is common, the major evolutionary grouping of assemblages should be segregated geographically. Thus, we would predict assemblages from South America (which was physical isolated through the Cenozoic) to constitute one group and assemblages from North and Central America another.

To test the relative importance of phylogenetic niche conservatism versus dispersal limitation, we analyzed data from *∼* 10, 000 tree assemblages with a new temporally-calibrated, genus-level phylogeny that includes 1,358 genera (*∼* 90% of tree genera sampled per assemblage). We assessed similarity in lineage composition among assemblages using clustering analyses and ordinations based on shared phylogenetic branch length. Next, we identified the indicator lineages for each major group in the clustering analysis and explored the geographic and environmental correlates of the distribution of the main evolutionary clusters. Finally, we estimated the unique evolutionary diversity (i.e. sum of phylogenetic branches of lineages restricted to individual groups) versus shared evolutionary diversity (i.e. sum of shared phylogenetic branches) across evolutionary groups (for details see SM).

We show that the evolutionary lineage composition of American tree assemblages is structured primarily by phylogenetic niche conservatism. The two principal groups have a tropics-extratropics structure (Fig. 1). The extratropical group is not geographically segregated, but includes temperate tree assemblages from North America and southern South America, connected by a high-elevation corridor in low latitudes (Fig. 1 a,b). The tropics-extratropics structure of tree evolutionary diversity shows a strong correspondence (97% match, Fig. S1) with the absence vs. occurrence of freezing temperatures within a typical year (see Fig. 1 c,d). We observe that most evolutionary diversity, measured as summed phylogenetic branch length, occurs within the tropics, but that there is unique evolutionary diversity restricted to the extratropics (*∼* 10% of the total, Fig. 2b, S3a). Ordination and indicator clade analyses revealed that the tropics-extratropics segregation is associated with the distribution of specific clades, such as the Fagales, which includes the oaks (*Quercus*), beeches (*Fagus*), coihues (*Nothofagus*) and their relatives (Fig. 3, Table S1, S2).

**Figure 1.**
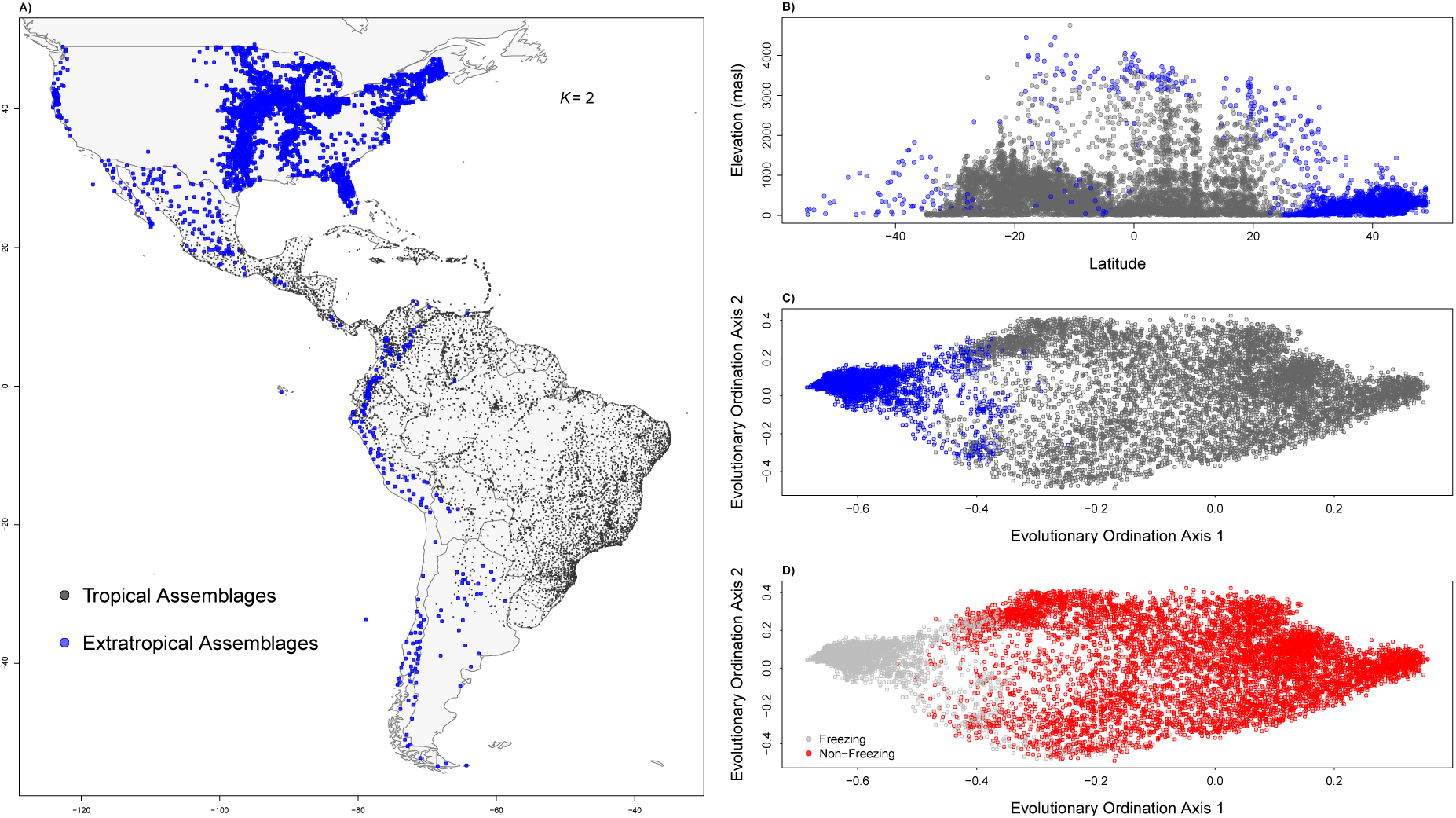
The geographic, evolutionary and environmental relationships of the two principal evolutionary groups (from *K*=2 clustering analysis). **A)** Geographic distribution of angiosperm tree assemblages and their affiliation with either the tropical (n = 7145) or extratropical (n = 2792) evolutionary group; **B)** Distribution of assemblages over elevation and latitude showing that the extratropical group is largely restricted to high elevations at low latitudes; **C & D)** Distribution of assemblages over the first two axes of an ordination based on evolutionary composition with assemblages in **C** colored according to group affiliation and in **D** as to whether or not they experience freezing temperatures in a regular year (from (*31*)).

**Figure 2.**
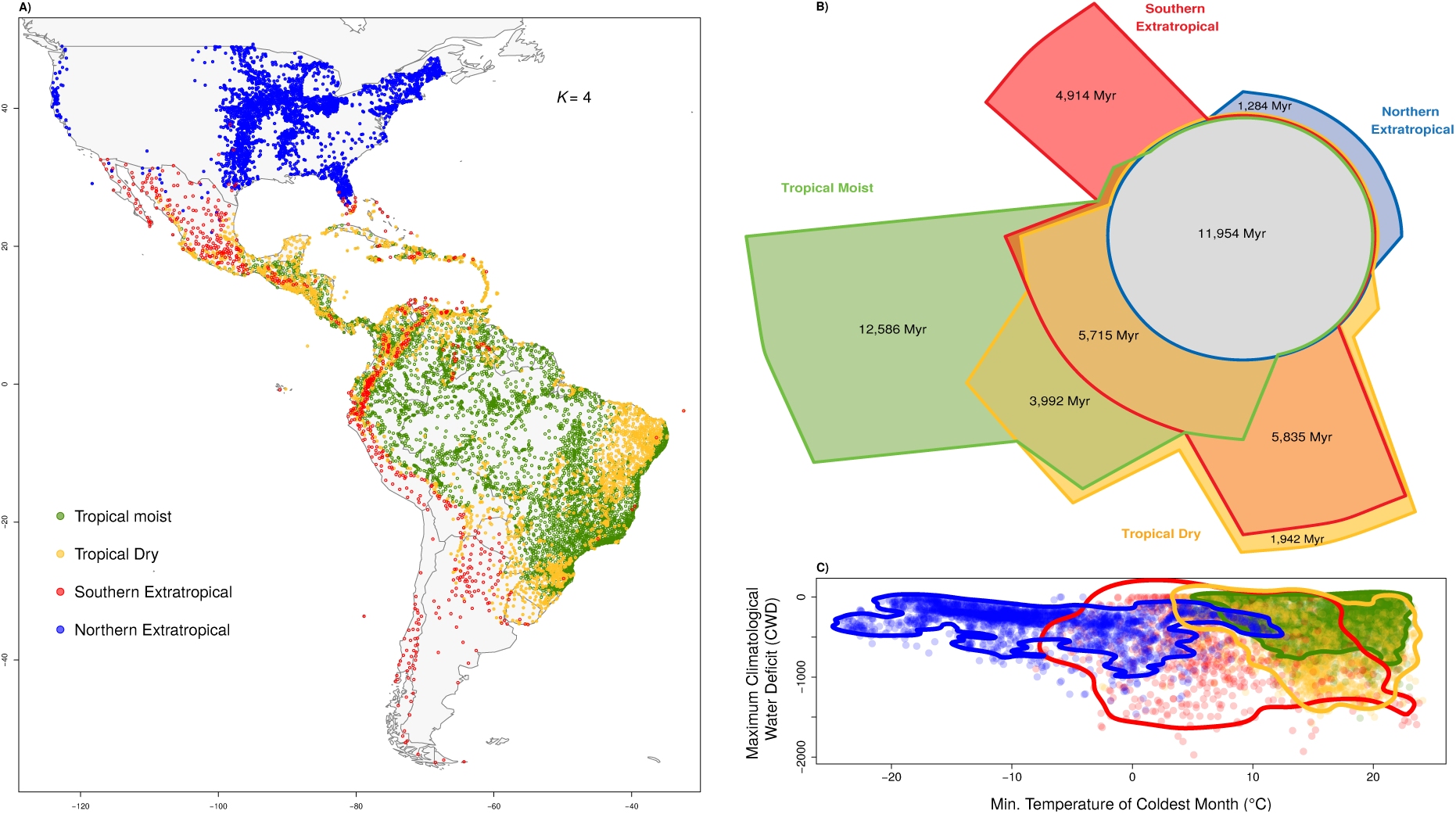
The geographic, evolutionary and environmental relationships among four evolutionary groups (from *K=*4 clustering analysis). **A)** Geographic distribution of angiosperm tree assemblages and their affiliation with one of the four evolutionary groups; **B)** Euler Diagram representing the amount of evolutionary history, quantified as phylogenetic diversity (in millions of years), restricted to each cluster versus that shared between clusters; **C)** Distribution of assemblages over extremes of temperature (minimum temperature of coldest month) and water availability (maximum climatological water deficit, CWD). Lines represent the 95th quantile of the density of points for each group.

**Figure 3.**
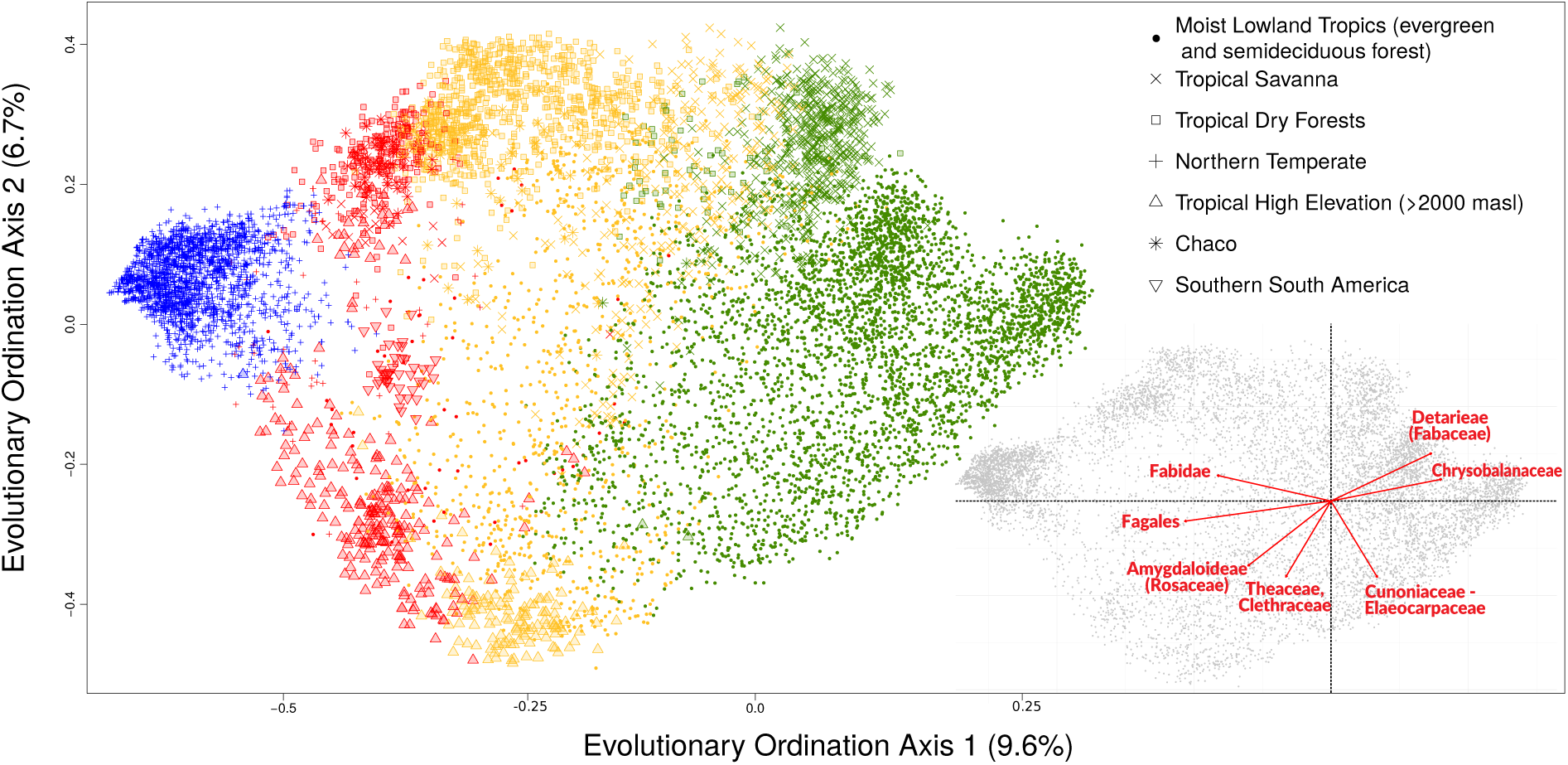
Phylogenetic ordination of tree assemblages based on their evolutionary lineage com- position. Colors in the main plot represent the groups from *K=*4 clustering analyses and the different symbols represent major vegetation formations. The subset plot shows the clades most strongly associated with the first two axes of the evolutionary ordination.

Cluster analyses of *K=*3 and *K=*4 groups are also supported as additional informative splits (Fig. S2), and each of their major groups capture substantial unique evolutionary diversity (Fig. 2b, Fig. S3, Table S2). In *K=*3, the main extratropical cluster grouped assemblages from North America and extreme southern South America, while the remaining assemblages from temperate southern South America and the Andean tropics grouped with assemblages from the arid or semiarid tropics and subtropics (Fig. S4). The third group was formed by the moist tropics (Fig. S4). For *K=*4, the extratropics were split into a largely temperate North American group and a second group that joins subtropical sites in South and Central America with southern temperate forests and high elevation sites in the Andes (Fig. 2a). In the tropics there is one group uniting assemblages found in ever-moist and warm conditions, and a second one of assemblages that extend into drier areas (Fig. 2c), including most tropical dry forest assemblages (Fig. 2a; Table S3). Hereafter, we refer to the four groups of assemblages in *K=*4 as the Northern Extratropical, Southern Extratropical, Tropical Moist and Tropical Dry groups.

Our results demonstrate that the tropics-extratropics evolutionary structure of tree diversity is principally associated with the environmental threshold of the presence versus absence of freezing temperatures (Fig. 1a,b, S1). This pattern is consistent with evidence documenting that only angiosperm lineages that were able to evolve traits to avoid freezing-induced embolism radiated into high latitudes (*12*). In addition, we found that a unique, sizeable portion of the total evolutionary diversity of angiosperm trees is restricted to extratropical environments, as the fossil record corroborates (*13, 14*). Collectively, this evidence suggests that the phylogenetic conservatism of lineages from the extratropics has a major relevance for the diversification of angiosperm trees in the Americas. Kerkhoff *et al.* (*15*) estimated that in the extratropical region (defined by them as areas north of 23°N and south of 23°S) angiosperm ancestors produced extratropical descendants at least 90% of the time. Considering that some areas subjected to regular freezing at high elevations in equatorial latitudes may be better classified as part of the extratropics, as demonstrated here by our results, extratropical phylogenetic conservatism could even be greater than found by Kerkhoff *et al.* (*15*).

We suggest that the extratropical conservatism has a major importance in the biogeography of the Americas. The relatively recent uplift of the Andes would have created novel environments, with regular freezing temperatures, at low latitudes, allowing extratropiocal lineages to move from both north and south to equatorial latitudes (*16, 17*). Fossil pollen demonstrates the arrival in the northern Andes of tree genera from temperate forests in the northern hemisphere, including *Juglans* (Juglandaceae), *Alnus* (Betulaceae) and *Quercus* (Fagaceae), at about 2.2 Ma, 1.0 Ma and 300 Ka respectively, and the arrival of southern genera, including *Weinmannia* (Cunoniaceae) and *Drymis* (Winteraceae), during the late Pliocene and Pleistocene (1.5–3.2 Ma) (*16, 18*). Likewise, phylogenetic evidence shows recent diversification in the Andes of lineages that seem to have originated in the extratropics, including *Lupinus* (Fabaceae) (*19*), Adoxaceae/Valerianceae (*20, 21*) and *Gunnera* (Gunneraceae) (*22*).

### Evolutionary Ordination Axis 1 (9.6%)

Our results also point to a moist versus dry evolutionary divide within the Neotropics. The Tropical Moist Group holds the greatest amount of evolutionary diversity, both overall and unique to it, despite occupying the most restricted extent of climatic space of any of the *K=*4 groups (Fig. 2 b,c). The Tropical Dry Group, in contrast, extends across a broader climatic space, but holds less evolutionary diversity (Fig. 2 b,c). This asymmetry in the accumulation of diversity may reflect phylogenetic conservatism for a putatively moist and hot ancestral angiosperm niche (*23*), or could result from a favorable environment that can be occupied by any angiosperm lineage, even those that also occur in cooler or drier conditions (*24, 25*). Regardless, the similarity in the lineage composition of the extensive but discontinuously distributed tropical dry forests (*11*) indicates their separate evolutionary history. Although tropical dry forest inhabiting taxa have often been described as more dispersal-limited than those from rain forests (e.g., *11*), dispersal over evolutionary time-scales seems to have been sufficient to maintain this floristic cohesion. Such evolutionary isolation of the dry forest flora has previously been suggested by studies in Fabaceae (*11, 26*), and is shown here to be evident at the evolutionary scale of all angiosperm tree species.

Our results also help to clarify the contentious evolutionary status of savanna and Chaco regions in the Neotropics. We find that the southern savannas (the Cerrado region of Brazil) are more evolutionary related to tropical moist forests than dry forests (Fig. 2 a, Fig. S4), as previously suggested (*26, 27*). However, northern tropical savannas (i.e., Llanos of Venezuela and Colombia and those in Central America) are split in their evolutionary linkages between the Tropical Moist and Tropical Dry groups (Fig. 3, Table S1). This may reflect the distinct ecology of many northern savannas (e.g., the Llanos are hydrological savannas; *28*) and suggest a divergent evolutionary history for northern and southern savannas. Our results may also help to resolve the debates around the evolutionary affinities of the Chaco (e.g., *29,30*), by showing that this geographically defined region houses a mix of extratropical and tropical lineages (Fig. 2). Furthermore, our analyses consistently point to evolutionary links between assemblages in seasonally dry and seasonally cold areas (Fig. 2, S4). For example, when we consider *K=*3 evolutionary groups, a single ‘dry and cool’ group coalesces, with the other two groups being the tropical moist forest group and a largely northern, extratropical group (Fig. S4).

We show that the evolutionary structure of tree diversity in the Americas is determined primarily by the presence versus absence of freezing temperatures, dividing tropical from extratropical regions. Within the tropics we find further subdivision among lineages experiencing moist versus seasonally-dry conditions. These findings clearly demonstrate that phylogenetic niche conservatism is the primary force organizing the diversification and, therefore, the biogeography of angiosperm trees. Tree species that can inhabit areas experiencing freezing temperatures and/or environments subjected to seasonal water stress belong to a restricted set of phylogenetic lineages, which gives a unique evolutionary identity to extratropical forests and tropical dry forests in the Americas. While our study is restricted to New World trees, we suggest that plant biodiversity globally may be evolutionarily structured following a tropics-extratropics pattern, while diversity within the tropics may be structured primarily around a moist-dry pattern. These findings advocate strongly for integrating the concept of extratropical conservatism and tropical-dry conservatism into our understanding of global macroevolutionary trends and biogeographic patterns.

## Materials and Methods

### Tree assemblage dataset

Our tree assemblage dataset was derived by combining the NeoTropTree (NTT) database (*32*) with selected plots from the Forest Inventory and Analysis (FIA) Program of the U.S. Forest Service (*33*), accessed on July 17th, 2018 via the BIEN package (*34*). We excluded from the latter any sites that had less than five angiosperm genera. Sites in the NTT database are dened by a single vegetation type within a circular area of 5-km radius and contains records of tree and tree-like species, i.e., freestanding plants with stems that can reach over 3m in height (see www.neotroptree.info and (*35*) for details). Each FIA plot samples trees that are *≥* 12.7 cm diameter at breast height (dbh) in four subplots (each being 168.3 m2) that are 36.6 m apart. We aggregated plots from the FIA dataset within 10 km diameter areas, to parallel the spatial structure of the NTT database. This procedure produced a total dataset of 9,937 tree assemblages distributed across major environmental and geographic gradients in the Americas.

### Genera phylogenetic tree

We obtained sequences of the *rbc*L and *mat*K plastid gene for 1,358 angiosperm tree genera, from Genbank (www.ncbi.nlm.nih.gov/genbank/), building on previous large-scale phylogenetic efforts for angiosperm trees in the Neotropics (*36, 37*). Sequences were aligned using the MAFFT software (*38*). ‘Ragged ends’ of sequences that were missing data for most genera were manually deleted from the alignment.

We estimated a maximum likelihood phylogeny for the genera in the RAxML v8.0.0 software (*39*), on the CIPRES web server (www.phylo.org). We constrained the tree to follow the order-level phylogeny in Gastauer *et al.* (2017) (*40*), which is based on the topology proposed by the Angiosperm Phylogeny Group IV. We concatenated the two chloroplast markers following a General Time Reversible (GTR) + Gamma (G) model of sequence evolution. We included sequences of *Nymphaea alba* (Nymphaeaceae) as an outgroup.

We temporally calibrated the maximum likelihood phylogeny using the software treePL (*41*). We implemented age constraints for 320 internal nodes (family-level or higher, from (*42*)) and for 123 genera stem nodes (based on ages from a literature survey, Table S4). The rate smoothing parameter (lambda) was set to 10 based on a cross-validation procedure. The final dated tree can be found in Supplementary Information.

### Phylogenetic distance analysis and clustering

We used the one complement of the Phylo-Sorensen Index (i.e., 1 – Phylo-Sorensen) to build a matrix of phylogenetic dissimilarities between plots based on genera presence-absence data. The Phylo-Sorensen index sums the total branch length of shared clades between sites (*43*) relative to the sum of branch lengths of both sites:

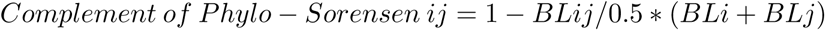

where BLij is the sum of branch lengths shared between plots i and j, and BLi and BLj are the sum of branch length of tips within plots i and j, respectively. Thus, if all branches are shared between two plots, the dissimilarity measure takes on a value of 0. If no branches are shared between plots (i.e. the plots comprise two reciprocally monophyletic clades), the dissimilarity measure will take on a value of 1. This metric was estimated using the phylosor.query() function in the PhyloMeasures (*44*) package for R.

We used *K-means* clustering to explore the main groups, in terms of (dis)similarity in the tree assemblage dataset, according to the Phylo-Sorensen dissimilarity measures. The *K-means* clustering algorithm requires the number of clusters (*K*) to be specified in advance. In order to estimate the best value for *K*, the optimal number of clusters to parsimoniously explain the variance in the dataset, we used the Elbow Method and an approach based on the average Silhouette width (Fig. S2). Based on these results, we selected *K=*2 (Fig. 1), *K=*3 (Fig. S5) and *K=*4 (Fig. 2) for further analysis and interpretation. No geographic or environmental data were used to inform the clustering analyses. The K-means clustering was carried out with the kmeans() function in base R (R Core Development Team, 2016). We assessed the robustness of the *K-means* clustering results using a silhouette analysis with functions in the “cluster” package (*45*). In order to assess variation in group fidelity, we classified individual sites as to whether the silhouette widths were larger or smaller than 0.2. In this way, we could detect areas of geographic, environmental and compositional space where clustering results were strongly or weakly supported.

In addition, we performed an evolutionary ordination of tree assemblages based on their phylogenetic lineage composition, following protocols developed by Pavoine (2016) (*46*). We specifically used an evolutionary PCA, implemented with the evopca() function in the “adiv” package (*47*), with a Hellinger transformation of the genus by site matrix, as this is a powerful approach to detect phylogenetic patterns along gradients, while also allowing positioning of sites and clades in an ordination space (*46*). The first two axes explained 9.6% and 6.7% of the variation in the data, with subsequent axes each explaining *<*5.5%.

### Correspondence between clustering results and environmental variables

We tested the correlation between our *K=2* clustering result and eight different delimitations of the tropics, as per Feeley and Stroud (2018) (*31*). These delimitations were: C1) all areas between 23.4°S and 23.4°N; C2) all areas with a net positive energy balance; C3) all areas where mean annual temperature does not co-vary with latitude; C4) all areas where temperatures do not go below freezing in a typical year; C5) all areas where the mean monthly temperature is never less than 18°C; C6) all areas where the mean annual “biotemperature” *≥* 24 °C; C7) all areas where the annual range of temperature is less than the average daily temperature range; C8) all areas where precipitation seasonality exceeds temperature seasonality. We calculated the correspondence between our binary clustering (i.e. tropical vs. extratropical) and each of these delimitations as the proportion of sites where the delimitations matched.

To assess the environmental space occupied by different groups from our clustering analyses, we obtained estimates of mean annual temperature, mean annual precipitation and minimum temperature of the coldest month from the Worldclim dataset (*48*) and Maximum Climatological Water Deficit (CWD) from Chave *et al.* (2014) (*49*). We estimated the density of the distribution of sites in the environmental space using ellipses containing 95% of the sites with the kde() function from *“ks”* package (*50*).

### Shared versus Unique “Phylogenetic Diversity” (PD)

As the Phylo-Sorensen estimation of evolutionary (dis)similarity cannot distinguish variation associated to differences in total phylogenetic diversity (PD), or phylogenetic richness versus variation associated to phylogenetic turnover per se, we measured the shared and unique PD associated with each group for the *K=*2, *K=*3 and *K=*4 clustering analyses. First, we estimated the association of genera with each group by an indicator species analysis following de Caceres *et al.* (2009) (*51*). Specifically, we used the multipatt() function in the R Packages indicspecies (*52*) to allow genera to be associated with more than one group (when *K >* 2). The output of the *multipatt ()* function includes the stat index, which is a function of the specificity (the probability that a surveyed site belongs to the target site group given the fact that the genus has been found) and fidelity (the probability of finding the genus in sites belonging to the given site group). We constructed pruned phylogenies including those genera with specificity greater than 0.6 to a group, or combination of groups, to estimate the total PD found in each group or combination of groups. Then, we subtracted these totals from the total for the complete, unpruned phylogeny to determine the amount of phylogenetic diversity restricted to each group, or combination of groups. Finally, we estimated the PD shared across all groups as that which was not restricted to any particular group or any combination of groups. We fit these different PD totals as areas in a Euler diagram with the euler() function in the *“eulerr”* package (*53*) for the *K=*2 and *K=*3 clustering, and with the Venn() fuction in the *“venn”* package (*54*) for the *K=*4 clustering.

### Indicator lineages for clusters

In order to further characterise the composition of the evolutionary groups, we conducted an indicator analysis to determine the clades most strongly associated with each group. We created a site x node matrix (see function used in Appendix 1), which consists of a presence/absence matrix for each internal node in the phylogeny and ran an indicator analysis for the nodes. We selected the highest-level, independent (i.e. non-nested) nodes with the highest stat values to present in Tables S1 and S2. The indicator node analysis was carried out with function *multipatt()* in the R Package indicspecies (*52*).

## Supplementary Materials

**Table S1.**
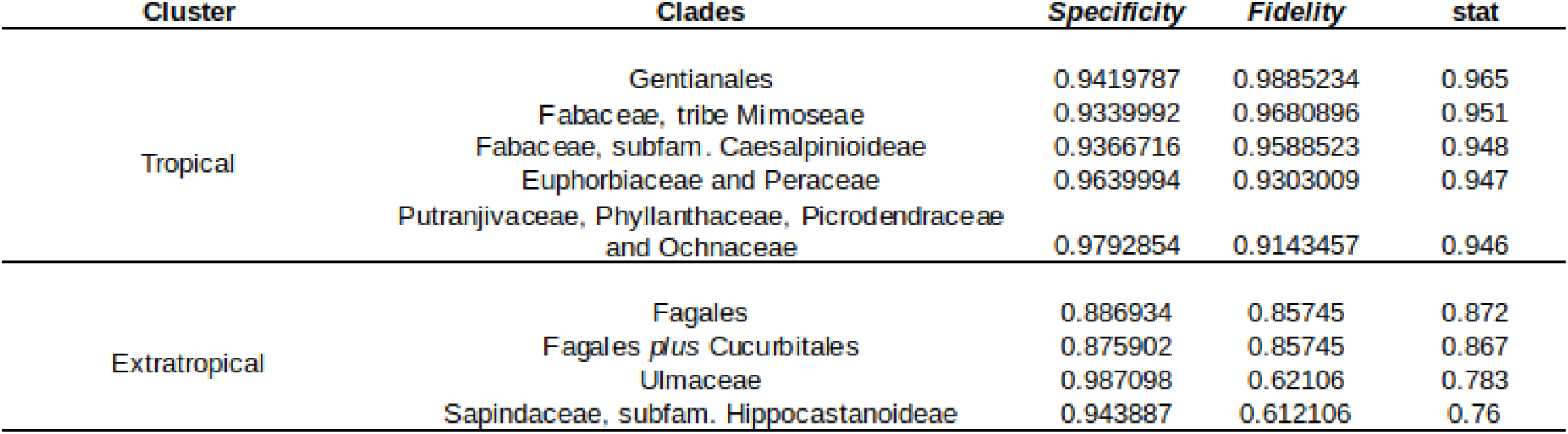
Indicator clades for *K*=2 groups. Specificity, fidelity and indicator statistic (stat) of internal nodes associated for the top nodes with the highest indicator statistic. Clades names are based on their taxonomic composition.

| Cluster | Clades | Specificity | Fidelity | stat |
| --- | --- | --- | --- | --- |
| Tropical | Gentianales | 0.9419787 | 0.9885234 | 0.965 |
|  | Fabaceae, tribe Mimoseae | 0.9339992 | 0.9680896 | 0.951 |
|  | Fabaceae, subfam. Caesalpinioideae | 0.9366716 | 0.9588523 | 0.948 |
|  | Euphorbiaceae and Peraceae | 0.9639994 | 0.9303009 | 0.947 |
|  | Putranjivaceae, Phyllanthaceae, Picrodendraceae and Ochnaceae | 0.9792854 | 0.9143457 | 0.946 |
| Extratropical | Fagales | 0.886934 | 0.85745 | 0.872 |
|  | Fagales *plus* Cucurbitales | 0.875902 | 0.85745 | 0.867 |
|  | Ulmaceae | 0.987098 | 0.62106 | 0.783 |
|  | Sapindaceae, subfam. Hippocastanoideae | 0.943887 | 0.612106 | 0.76 |

**Table S2.** Indicator clades for *K*=4 groups. Specificity, fidelity and indicator statistic (stat) of internal nodes associated for the top nodes with the highest indicator statistic. Clade names are based on their taxonomic composition.

| Cluster | Clades | Specificity | Fidelity | stat |
| --- | --- | --- | --- | --- |
| Tropical Moist | *Xylopia*, *Fusaea* and *Duguetia* (Annonaceae) | 0.849969 | 0.700198 | 0.771 |
|  | *Couepia* and *Hirtella* (Chrysobalanaceae) | 0.757572 | 0.711399 | 0.734 |
|  | Burseraceae, tribes Protieae and Canarieae | 0.794147 | 0.678014 | 0.734 |
|  | Ochnaceae | 0.746723 | 0.706128 | 0.726 |
|  | Myristicaceae | 0.851393 | 0.591917 | 0.71 |
|  | Calophyllaceae | 0.7516 | 0.668351 | 0.709 |
| Tropical Dry | Bignoniaceae, tribes Bignonieae and Tecomeae | 0.778906 | 0.117489 | 0.303 |
|  | *Lasiocarpus* and *Ptilochaeta* (Malpighiaceae) | 0.853331 | 0.072646 | 0.249 |
|  | Cactaceae, tribe Trichocereae | 0.864368 | 0.047982 | 0.204 |
| Southern Extratropical | *Prosopis*, *Piptadeniopsis* (Fabaceae, tribe Mimoseae) | 0.733744 | 0.402597 | 0.544 |
|  | Fabaceae, tribe Caesalpinieae | 0.670481 | 0.301587 | 0.45 |
|  | *Vallea* and *Aristotelia* (Elaeocarpaceae) | 0.854485 | 0.223665 | 0.437 |
|  | Cactaceae, tribes Pachycereae and Notocactea | 0.732889 | 0.248196 | 0.426 |
|  | Scrophulariaceae | 0.597146 | 0.30303 | 0.425 |
| Northern Extratropical | Sapindaceae, subfam. Hippocastanoideae | 0.83986 | 0.69118 | 0.762 |
|  | Ulmaceae | 0.75668 | 0.70459 | 0.73 |
|  | Oleaceae, tribe Oleae | 0.80497 | 0.62332 | 0.708 |
|  | Juglandaceae | 0.76212 | 0.53677 | 0.64 |

**Table S3.** Affiliation of principal vegetation formations in the tropics with the two main tropical groups from the K=4 clustering analysis. Vegetation formations were taken from the NeoTropTree dataset, which categorises formations first based on physiognomy (savanna vs. forest) and then segregates the forests based on phenology. Following (*35*) and (*54*), we consider deciduous tropical forests to represent the tropical dry forest biome, while semideciduous forests are more related floristically to the tropical moist forest biome. Semideciduous forests share many tree species with evergreen forests and relatively few with more fully deciduous forests (*35, 54*). We further divided the savannas based on geography, as our analyses showed evident differences in group affiliation between savannas in the Cerrado Domain of Brazil versus those further north (i.e. Llanos of Venezuela and Colombia and those in Central America).

|  | Tropical dry | Tropical moist |
| --- | --- | --- |
| Evergreen Forests | 15% (501) | 85% (2948) |
| Semideciduous Forests | 10% (167) | 90% (1530) |
| Deciduous Forests | 75% (868) | 25% (285) |
| Southern Savannas (Cerrado) | 8% (56) | 92% (657) |
| Northern Savannas | 54% (65) | 46% (56) |

**Table S4.** Stem ages for genera nodes used to callibrate the phylogenetic tree, and the reference of their source. References (*55–94*).

| Genus | stem age (Myr) | Reference |
| --- | --- | --- |
| 1 *Acer* | 60 | Renner et al 2008, Systematic Biology |
| 2 *Acioa* | 19.1 | Bardon et al 2016, American Journal of Botany |
| 3 *Aesculus* | 65 | Harris et al 2009, Taxon |
| 4 *Anaxagorea* | 90.44 | Baker et al. 2014, Ecology Letters (and references herein) |
| 5 *Andira* | 17.51 | Baker et al. 2014, Ecology Letters (and references herein) |
| 6 *Antiaris* | 34 | Gardner et al 2017, Molecular Phylogenetics and Evolution |
| 7 *Aphananthe* | 71.5 | Yang et al 2007, PLOS ONE |
| 8 *Aphanocalyx* | 46 | Bruneau et al 2008, Botany |
| 9 *Artocarpus* | 51 | Rockinger et al 2017, BMC Evolutionary Biology |
| 10 *Atuna* | 20.5 | Bardon et al 2016, American Journal of Botany |
| 11 *Avicennia* | 70.09 | Tripp & McDade 2014, Systematic Biology |
| 12 *Bagassa* | 67 | Gardner et al 2017, Molecular Phylogenetics and Evolution |
| 13 *Bocageopsis* | 5.98 | Baker et al. 2014, Ecology Letters (and references herein) |
| 14 *Brosimum* | 48 | Baker et al. 2014, Ecology Letters (and references herein) |
| 15 *Caesalpinia* | 48.3 | Bruneau et al 2008, Botany |
| 16 *Carapa* | 29.5 | Baker et al. 2014, Ecology Letters (and references herein) |
| 17 *Cassia* | 45 | Bruneau et al 2008, Botany |
| 18 *Castilla* | 22 | Baker et al. 2014, Ecology Letters (and references herein) |
| 19 *Cecropia* | 44 | Baker et al. 2014, Ecology Letters (and references herein) |
| 20 *Cedrela //toona* | 48.4 | Muellner et al 2010, American Journal of Botany |
| 21 *Cedrelopsis* | 18.94 | Appelhans et al 2012, Journal of Biogeography |
| 22 *Centroplacus* | 69 | Cai et al 2016, PlosONE |
| 23 *Cercis* | 47.3 | Bruneau et al 2008, Botany |
| 24 *Chrysobalanus* | 24.2 | Bardon et al 2016, American Journal of Botany |
| 25 *Cissus* | 67.99 | Rodrigues et al 2014, Taxon |
| 26 *Clarisia* | 70 | Gardner et al 2017, Molecular Phylogenetics and Evolution |
| 27 *Coceveiba* | 72 | Baker et al. 2014, Ecology Letters (and references herein) |
| 28 *Cornus* | 74.03 | Xiang et al 2011, Molecular Phylogenetics and Evolution |
| 29 *Couepia* | 21.6 | Bardon et al 2016, American Journal of Botany |
| 30 *Crudia* | 45 | Bruneau et al 2008, Botany |
| 31 *Cylicomorpha* | 35.5 | Antunes & Renner 2012, Molecular Phylogenetics and Evolution |
| 32 *Cynometra* | 12.93 | Baker et al. 2014, Ecology Letters (and references herein) |
| 33 *Dacryodes* | 38 | Baker et al. 2014, Ecology Letters (and references herein) |
| 34 *Dactyladenia* | 15.9 | Bardon et al 2016, American Journal of Botany |
| 35 *Dactylocladus* | 39 | Moyle 2004, Evolution |
| 36 *Dialium* | 10.9 | Baker et al. 2014, Ecology Letters (and references herein) |
| 37 *Dicymbe* | 12 | Baker et al. 2014, Ecology Letters (and references herein) |
| 38 *Diplotropis* | 20.27 | Baker et al. 2014, Ecology Letters (and references herein) |
| 39 *Dipterix* | 26.44 | Baker et al. 2014, Ecology Letters (and references herein) |
| 40 *Dipterocarpus* | 47.7 | Heckenhauer et al 2017, Botanical Journal of the Linnean Society |
| 41 *Drimys* | 56.76 | Thomas et al 2014, Journal of Biogeography |
| 42 *Dryobalanops* | 43.3 | Heckenhauer et al 2017, Botanical Journal of the Linnean Society |
| 43 *Duguetia* | 30.64 | Baker et al. 2014, Ecology Letters (and references herein) |
| 44 *Dycorynia* | 10.9 | Baker et al. 2014, Ecology Letters (and references herein) |
| 45 *Embothrium* | 39.3 | Sauquet et al 2009, PNAS |
| 46 *Eperua* | 12.32 | Baker et al. 2014, Ecology Letters (and references herein) |
| 47 *Ficus* | 58 | Gardner et al 2017, Molecular Phylogenetics and Evolution |
| 48 *Froesia* | 39.4 | Schneider & Zizka 2017, Taxon |
| 49 *Fusaea* | 30.64 | Baker et al. 2014, Ecology Letters (and references herein) |
| 50 *Glochidion* | 31.51 | van Welzen et al 2015, Journal of Biogeography |
| 51 *Glycosmis* | 32.54 | Appelhans et al 2012, Journal of Biogeography |
| 52 *Guarea* | 14.8 | Baker et al. 2014, Ecology Letters (and references herein) |
| 53 *Guatteria* | 55.83 | Baker et al. 2014, Ecology Letters (and references herein) |
| 54 *Gyrocarpus* | 72 | Michalak et al 2010, Journal of Biogeography |
| 55 *Hakea* | 12.8 | Mast et al 2012, American Journal of Botany |
| 56 *Harrisoia* | 57.99 | Appelhans et al 2012, Journal of Biogeography |
| 57 *Helicostylis* | 28 | Baker et al. 2014, Ecology Letters (and references herein) |
| 58 *Hennecartia* | 15.58 | Renner et al 2010, Journal of Biogeography |
| 59 *Hernandia* | 76 | Michalak et al 2010, Journal of Biogeography |
| 60 *Hevea* | 85 | Baker et al. 2014, Ecology Letters (and references herein) |
| 61 *Hopea* | 21.6 | Heckenhauer et al 2017, Botanical Journal of the Linnean Society |
| 62 *Hymenaea* | 24 | Bruneau et al 2008, Botany |
| 63 *Inga* | 10 | Baker et al. 2014, Ecology Letters (and references herein) |
| 64 *Ipomoea* | 34.97 | Eserman et al 2014, American Journal of Botany |
| 65 *Iryanthera* | 19 | Baker et al. 2014, Ecology Letters (and references herein) |
| 66 *Jacaratia* | 27.5 | Antunes & Renner 2012, Molecular Phylogenetics and Evolution |
| 67 *Lacunaria* | 20.3 | Schneider & Zizka 2017, Taxon |
| 68 *Lomatia* | 70.8 | Milner et al 2015, Journal of Biogeography |
| 69 *Lonchocarpus* | 15.07 | Baker et al. 2014, Ecology Letters (and references herein) |
| 70 *Lonicera* | 43.37 | Bell & Donoghue 2005, American Journal of Botany |
| 71 *Maclura* | 85 | Gardner et al 2017, Molecular Phylogenetics and Evolution |
| 72 *Macrolobium* | 32 | Baker et al. 2014, Ecology Letters (and references herein) |
| 73 *Magnistipula* | 19 | Bardon et al 2016, American Journal of Botany |
| 74 *Malmea* | 19.99 | Baker et al. 2014, Ecology Letters (and references herein) |
| 75 *Manilkara* | 32 | Armstrong et al 2014, Frontiers in Genetics |
| 76 *Maranthes* | 20.5 | Bardon et al 2016, American Journal of Botany |
| 77 *Meliosma* | 67.34 | Yang et al 2018, Molecular Phylogenetics and Evolution |
| 78 *Mimusops* | 35 | Armstrong et al 2014, Frontiers in Genetics |
| 79 *Mouriri* | 90 | Renner 2004, American Journal of Botany |
| 80 *Myrtae tribe* | 58.96 | Vasconcelos et al 2017, Molecular Phylogenetics and Evolution |
| 81 *Neocarya* | 25.6 | Bardon et al 2016, American Journal of Botany |
| 82 *Ormosia* | 40.62 | Baker et al. 2014, Ecology Letters (and references herein) |
| 83 *Otoba* | 17 | Baker et al. 2014, Ecology Letters (and references herein) |
| 84 *Parashora* | 22.9 | Heckenhauer et al 2017, Botanical Journal of the Linnean Society |
| 85 *Parkia* | 45.5 | Baker et al. 2014, Ecology Letters (and references herein) |
| 86 *Peltogyne* | 28.8 | Baker et al. 2014, Ecology Letters (and references herein) |
| 87 *Persea* | 55.3 | Li et al 2018, American Journal of Botany |
| 88 *Peumus* | 55.66 | Renner et al 2010, Journal of Biogeography |
| 89 *Poecilanthe* | 40.99 | Baker et al. 2014, Ecology Letters (and references herein) |
| 90 *Poulsenia* | 22 | Baker et al. 2014, Ecology Letters (and references herein) |
| 91 *Pourouma* | 44 | Baker et al. 2014, Ecology Letters (and references herein) |
| 92 *Pradosia* | 47.5 | Terra-Araujo et al 2015, Molecular Phylogenetics and Evolution |
| 93 *Prosopis SA* | 28.96 | Catalano et al 2008, Botanical Journal of the Linnean Society |
| 94 *Protium* | 52.5 | Baker et al. 2014, Ecology Letters (and references herein) |
| 95 *Prunus* | 60.7 | Chin et al 2014, Molecular Phylogenetics and Evolution |
| 96 *Pseudolmedia* | 36 | Baker et al. 2014, Ecology Letters (and references herein) |
| 97 *Pseudowintera* | 45.18 | Thomas et al 2014, Journal of Biogeography |
| 98 *Pseudoxandra* | 15.09 | Baker et al. 2014, Ecology Letters (and references herein) |
| 99 *Pterocarpus* | 16.66 | Baker et al. 2014, Ecology Letters (and references herein) |
| 100 *Quiina* | 29 | Schneider & Zizka 2017, Taxon |
| 101 *Rhododendron* | 58 | Schwery et al 2015, New Phytologist |
| 102 *Richea* | 22.31 | Schwery et al 2015, New Phytologist |
| 103 *Sambucus* | 45.49 | Bell & Donoghue 2005, American Journal of Botany |
| 104 *Sideroxylon* | 74 | Smedmark & Anderberg 2007, American Journal of Botany |
| 105 *Slonaea* | 79 | Crayn et al 2006, American Journal of Botany |
| 106 *Sorocea* | 59 | Baker et al. 2014, Ecology Letters (and references herein) |
| 107 *Spathelia* | 19.21 | Appelhans et al 2012, Journal of Biogeography |
| 108 *Swartzia* | 45.96 | Baker et al. 2014, Ecology Letters (and references herein) |
| 109 *Tachigali* | 4.65 | Baker et al. 2014, Ecology Letters (and references herein) |
| 110 *Tasmania* | 69.98 | Thomas et al 2014, Journal of Biogeography |
| 111 *Tepualia* | 24.9 | Thornhill et al 2015, Molecular Phylogenetics and Evolution |
| 112 *Theobroma* | 11.6 | Richardson et al 2015, Front. Ecol. Evol. |
| 113 *Unonopsis* | 7.94 | Baker et al. 2014, Ecology Letters (and references herein) |
| 114 *Urophyllum* | 27.1 | Smedmark et al 2010, Journal of Biogeography |
| 115 *Vallea* | 48 | Crayn et al 2006, American Journal of Botany |
| 116 *Vateria* | 15.4 | Heckenhauer et al 2017, Botanical Journal of the Linnean Society |
| 117 *Vatica* | 18.3 | Heckenhauer et al 2017, Botanical Journal of the Linnean Society |
| 118 *Viburnum* | 71.18 | Bell & Donoghue 2005, American Journal of Botany |
| 119 *Virola* | 17 | Baker et al. 2014, Ecology Letters (and references herein) |
| 120 *Vitellariopsis* | 26 | Armstrong et al 2014, Frontiers in Genetics |
| 121 *Vouacapoua* | 48.69 | Baker et al. 2014, Ecology Letters (and references herein) |
| 122 *Xylopia* | 49.98 | Baker et al. 2014, Ecology Letters (and references herein) |
| 123 *Zygia* | 17.82 | Baker et al. 2014, Ecology Letters (and references herein) |

**Fig. S1.**
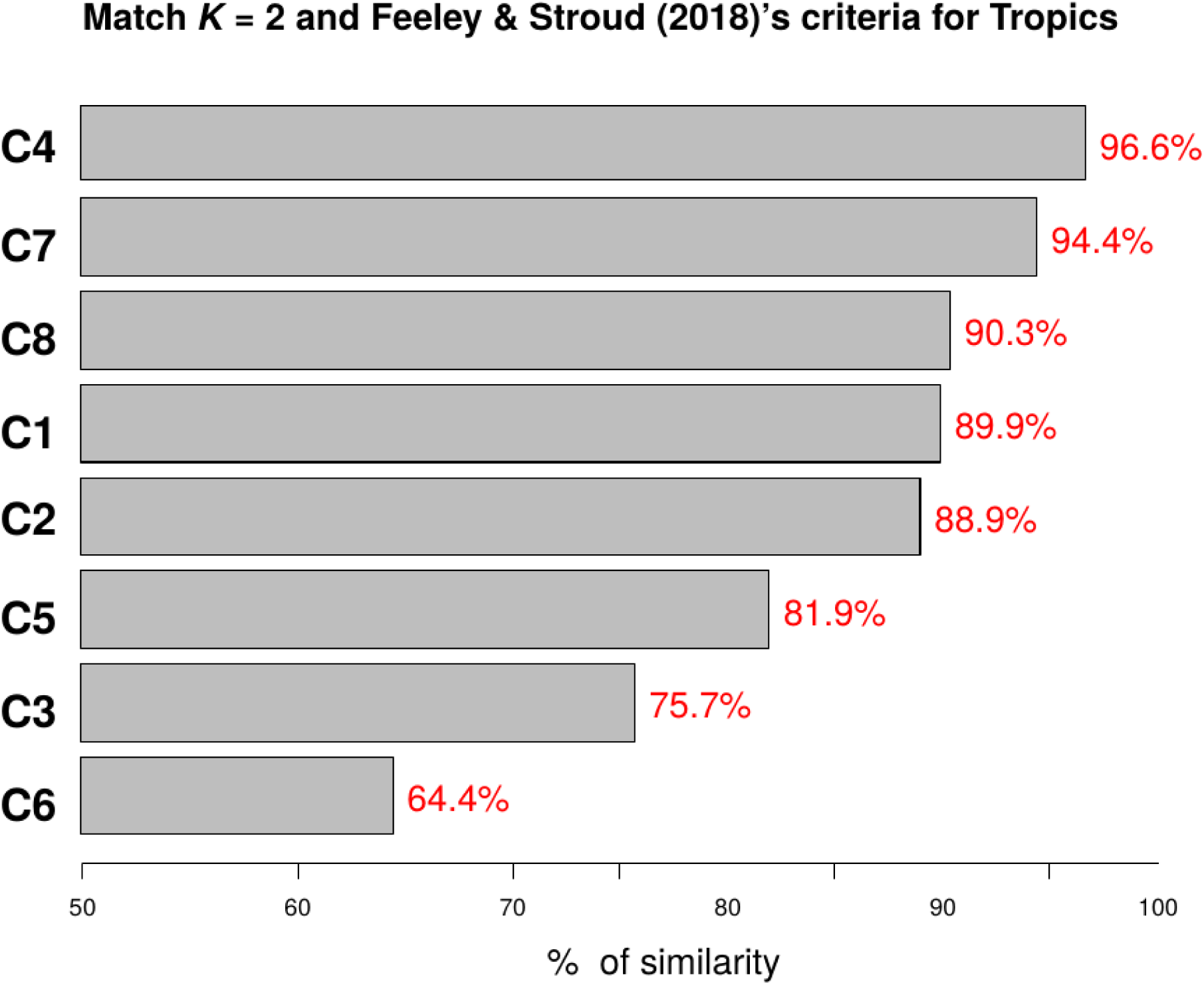
Match between tropics vs. extratropics groups from K=2 clustering and eight delimitations of the tropics following Feeley & Stroud [2018] (*31*): C1) all areas that occur between 23.4°S and 23.4°N; C2) all areas with a net positive energy balance; C3) all areas where mean annual temperature does not vary with latitude; C4) all areas where temperatures do not go below freezing in a typical year; C5) all areas where the mean monthly temperature is never less than 18°C; C6) all areas where the mean annual “biotemperature” ≥ 24°C; C7) all areas where the annual range of temperature is less than the average daily temperature range; C8) all areas where precipitation seasonality exceeds temperature seasonality.

**Fig. S2.**
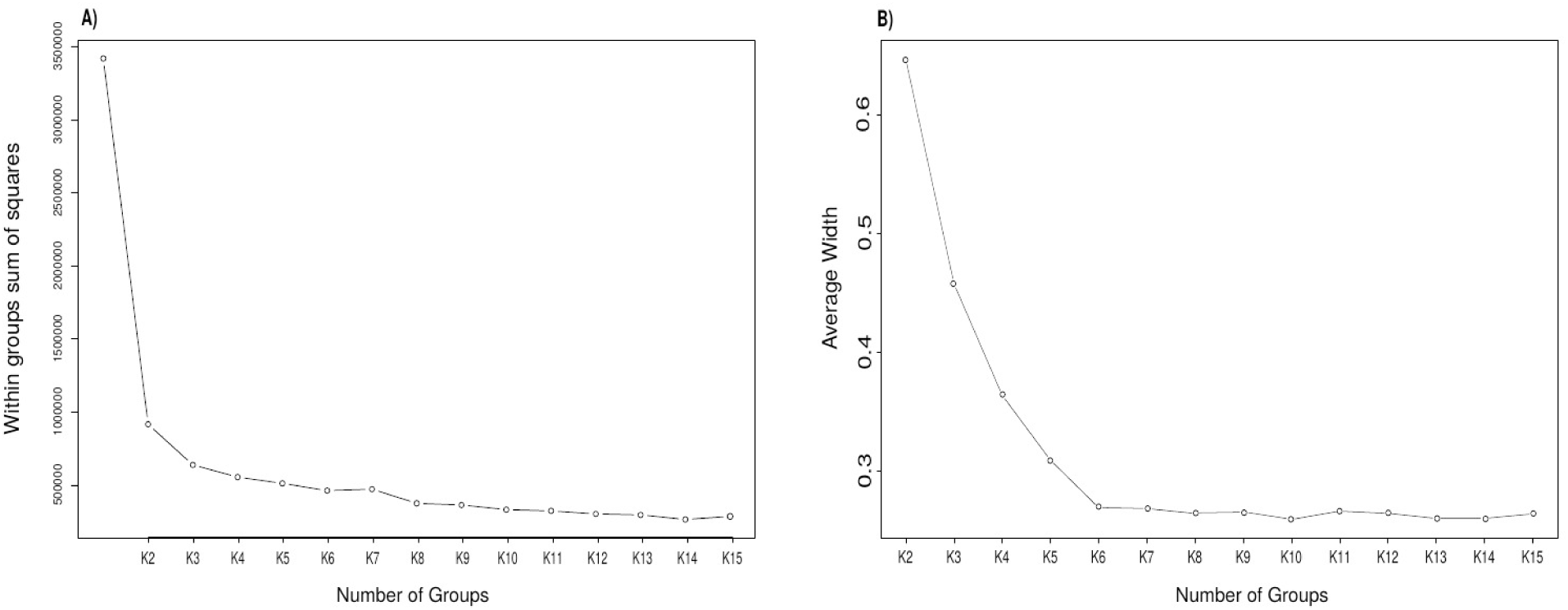
Selection of number of clusters. A) Elbow criterion, explained variance from clustering as a function of number of groups; B) Silhouette criterion, average silhouette width for each site as a function of number of groups.

**Fig. S3.**
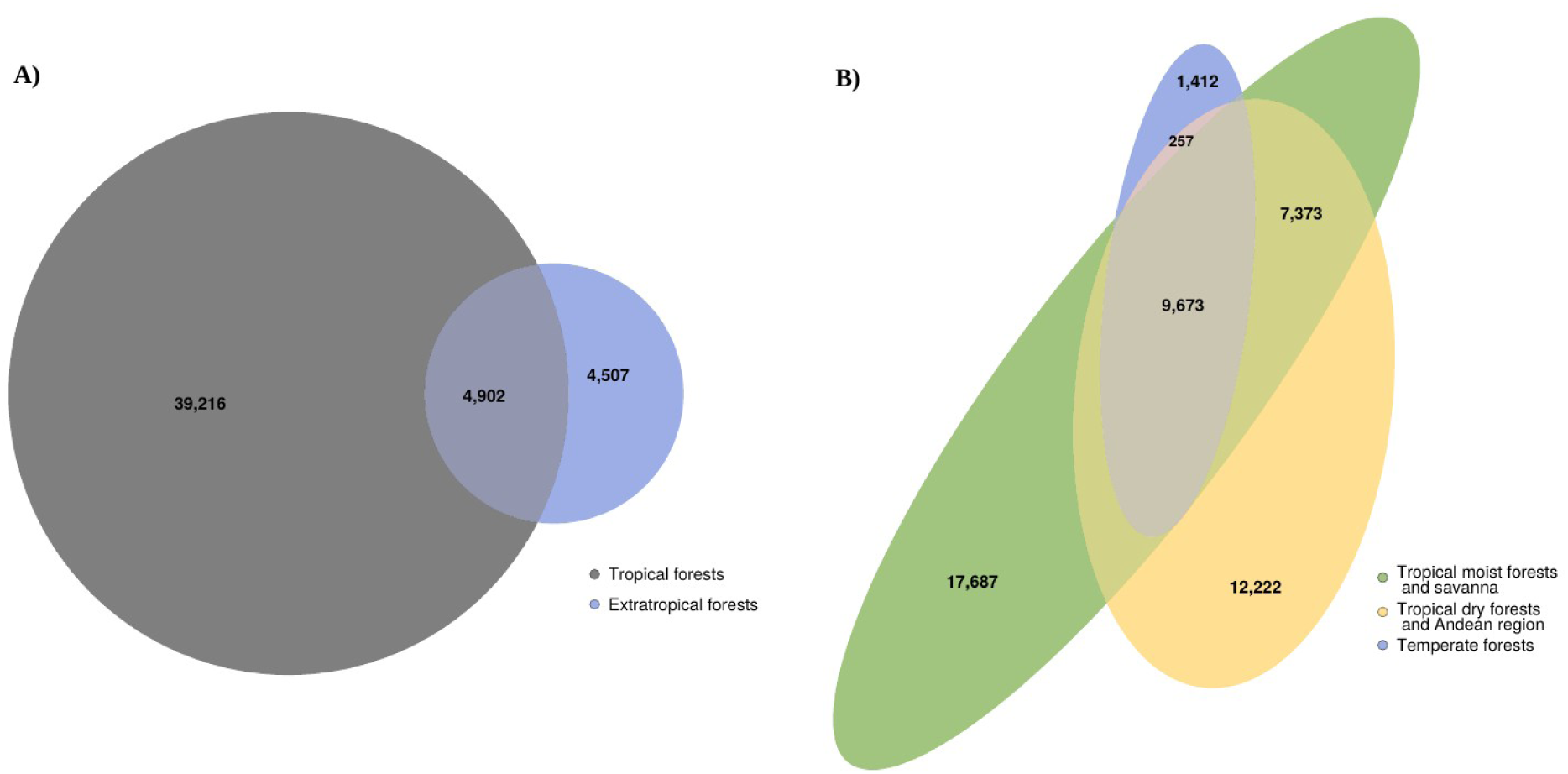
Shared versus unique Phylogenetic Diversity for *K*=2 and *K*=3 clustering analyses. Euler Diagrams showing the amount of unique phylogenetic diversity in each cluster and the phylogenetic diversity shared between clusters (in millions of years). A) *K*=2 clustering and B) *K*=3 clustering.

**Fig. S4.**
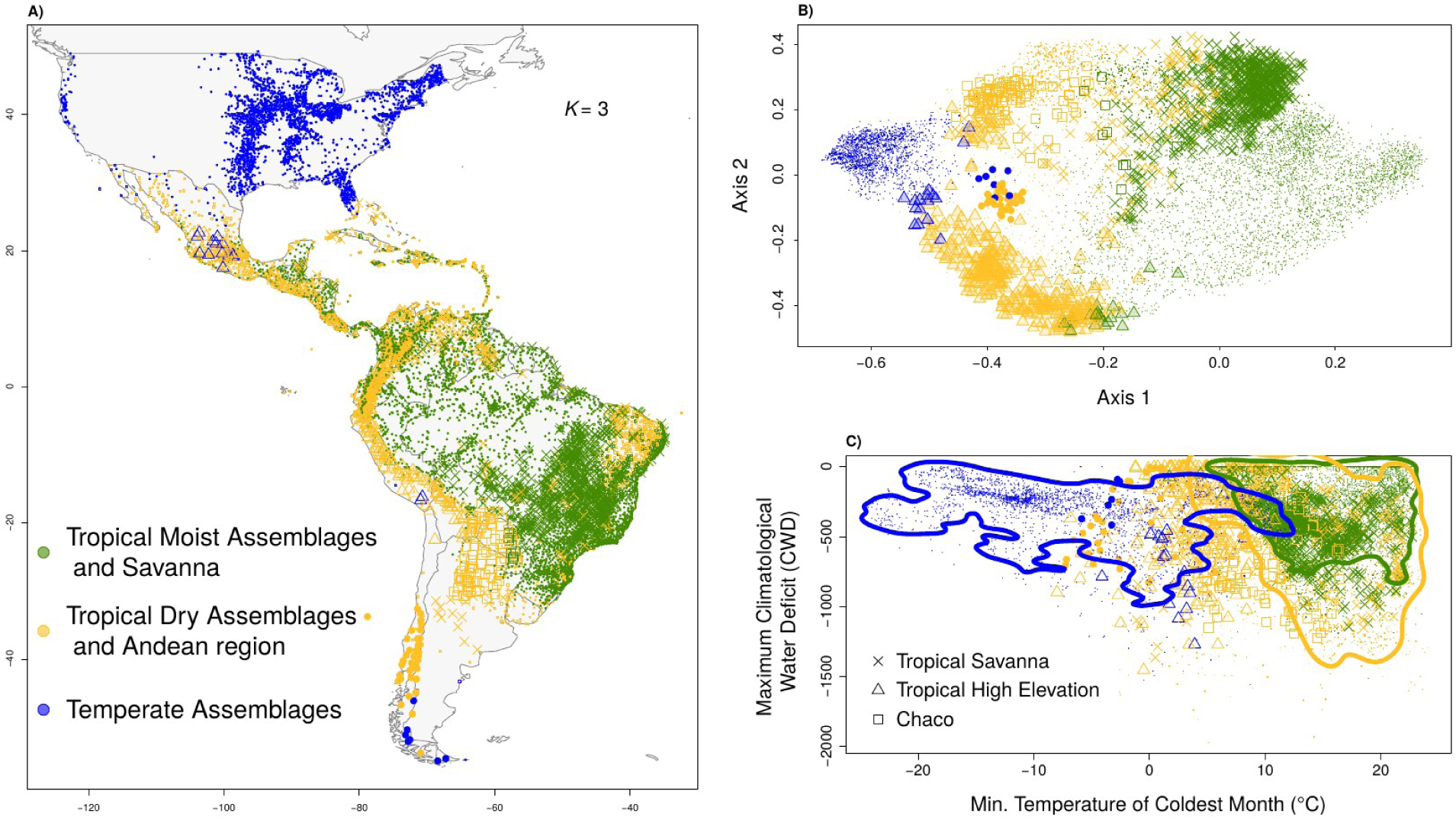
Clustering *K* = 3. A) Location of the 9937 angiosperm tree assemblages in three evolutionary groups; B) Ordination of tree assemblages based on evolutionary lineage composition; C) Maximum Climatological Water Deficit (CWD) versus minimum temperature of the coldest month. Lines represent the 95th quantile of the density of points for each group. In each panel, symbol type indicates major vegetation type.

**Fig. S5.**
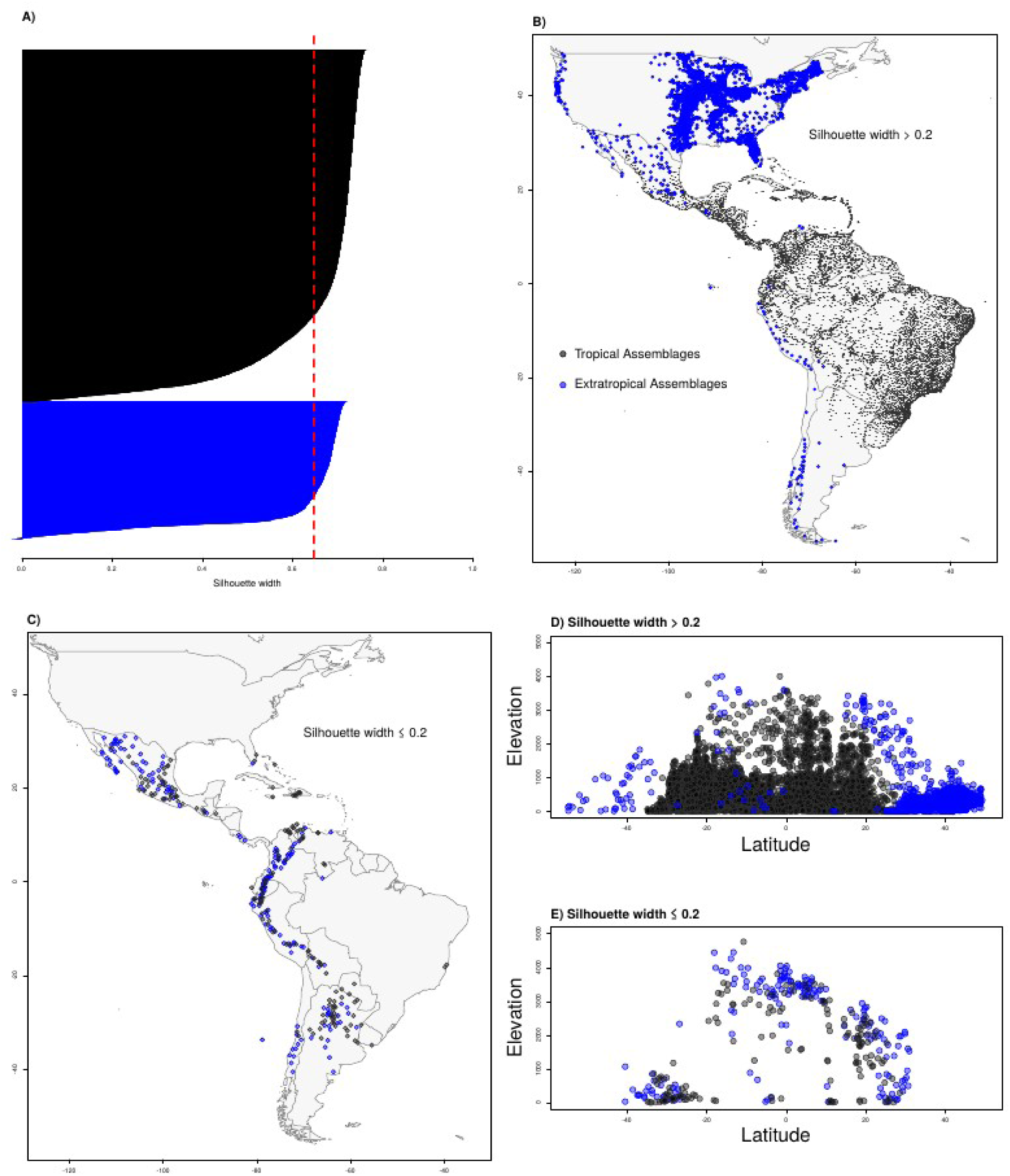
Clustering validation *K*= 2. A) Silhouette width for each assemblage in our dataset. The colors represent the clusters and the red dashed line the average silhouette width. B) shows the distribution of sites with silhouette > 0.2, which indicates that they are strongly associated with their given cluster. C) shows the geographic distribution of sites with silhouette width ≤ 0.2. D) shows sites with silhouette width > 0.2 in a latitude-elevation plot and E) shows sites with silhouette width ≤ 0.2 in a latitude-elevation plot.

**Fig. S6.**
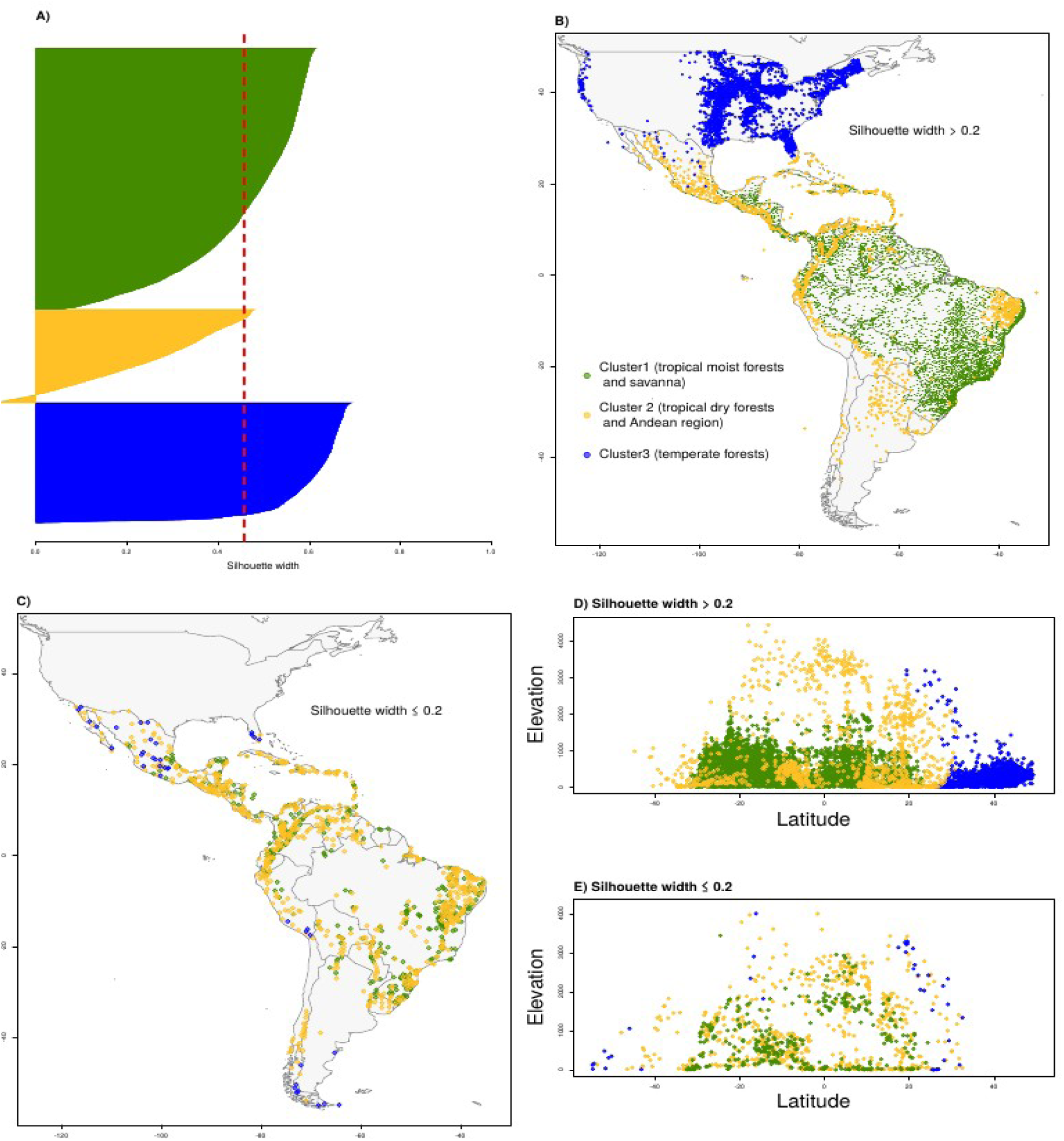
Clustering validation *K*= 3. A) Silhouette width for each assemblage in our dataset. The colors represent the clusters and the red dashed line the average silhouette width. B) shows the distribution of sites with silhouette width bigger than 0.2, which indicates that they are strongly associated with their given cluster. C) shows the geographic distribution of sites with silhouette width < 0.2. D) shows sites with silhouette width > 0.2 in a latitude-elevation plot and E) shows sites with silhouette width < 0.2 in a latitude-elevation plot.

**Fig. S7.**
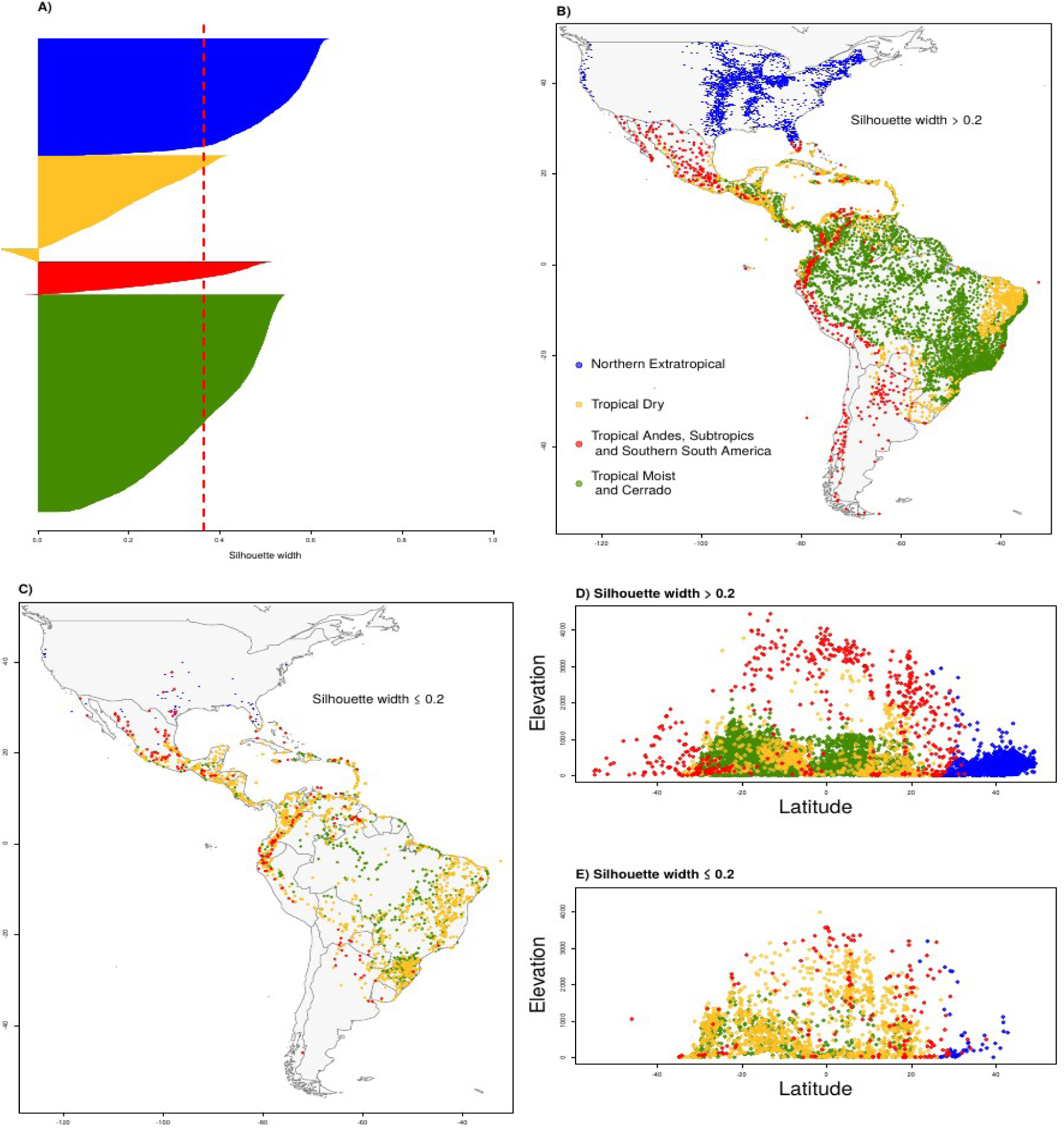
Clustering validation *K*= 4. A) Silhouette width for each assemblage in our dataset. The colors represent the clusters and the red dashed line the average silhouette width. B) shows the distribution of sites with silhouette width bigger than 0.2, which indicates that they are strongly associated with their givencluster. C) shows the geographic distribution of sites with silhouette width < 0.2. D) shows sites with silhouette width > 0.2 in a latitude-elevation plot and E) shows sites with silhouette width < 0.2 in a latitude-elevation plot.

